# KLF7 Promotes Preadipocyte Proliferation via Activation of the Akt Signaling Pathway by Cis-regulating CDKN3

**DOI:** 10.1101/2022.06.16.496506

**Authors:** Ziqiu Jia, Zhao Jin, Shuli Shao, Hu Xu, Wen Li, Mahmood Khan, Weiyu Wang, Weiwei Zhang, Yingning Sun

## Abstract

Krüppel-like transcription factor 7 **(KLF7)** promotes preadipocyte proliferation; however, its target gene in this process has not yet been identified. Using KLF7 ChIP-seq analysis, we previously showed that a KLF7-binding peak is present upstream of the cyclin-dependent kinase inhibitor 3 gene **(*CDKN*3)** in chicken preadipocytes. In the current study, we identified *CDKN*3 as a target gene of KLF7 that mediates the effects of KLF7 on preadipocyte proliferation. Furthermore, 5′-truncating mutation analysis showed that the minimal promoter was located between *CDKN*3 nt −160 and nt −7 (relative to the translation initiation codon ATG). KLF7 overexpression increased *CDKN*3 promoter activity in the DF-1 and immortalized chicken preadipocyte **(ICP1)** cell lines. Deletion of the putative binding site of KLF7 abolished the promotive effect of KLF7 overexpression on *CDKN*3 promoter activity. Moreover, *CDKN3*-knockdown and -overexpression assays revealed that CDKN3 enhanced ICP1 cell proliferation. Flow cytometry analysis showed that CDKN3 accelerated the G1/S transition. Further, we found that KLF7 promoted ICP1 cell proliferation via Akt phosphorylation by regulating CDKN3. Taken together, these results suggest that KLF7 promotes preadipocyte proliferation via activating the Akt signaling pathway by cis-regulating *CDKN*3, thus driving the G1/S transition.

## Introduction

In recent years, the incidences of type 2 diabetes, coronary heart disease, and other obesity-related diseases have increased rapidly, and these diseases have become a major public health problem [1]. Obesity is mainly caused by excessive fat deposition [2], which is associated with an increase in the number and size of adipocytes. Adipocyte number mainly attribute to preadipocyte proliferation, whereas adipocyte size attribute to preadipocyte differentiation [3]. Preadipocyte proliferation is a complex process controlled by a regulatory network of multiple factors, including transcription factors [4-6], cytokines [7], and non-coding RNAs [8, 9].

The Krüppel-like transcription factors (**KLFs**) are essential for lots of physiological and pathological processes [10, 11]. KLF7 is associated with obesity [12, 13], type 2 diabetes [14], and cardiovascular and cerebrovascular diseases [15, 16]. KLF7 inhibits preadipocyte differentiation in mammals and birds [4, 17], and promotes chicken preadipocyte proliferation [4]. In separated chicken stromal-vascular cells, KLF7-overexpressing cells exhibited an enhanced proliferation ability, especially, at 48 h and 120 h after transfection, when compared with the control cells [4]. The negative effect of KLF7 on preadipocyte differentiation is mainly achieved via the regulation of target genes such as *GATA*3 [18] and *HIF*1*A* [19]. However, the target gene of KLF7 in the process of promoting chicken preadipocyte proliferation remains unknown.

Chicken is an ideal model for studying adipogenesis, obesity, and adipose biology [20-22]. In our previous study using ChIP-seq analysis of chicken preadipocytes, we found a KLF7-binding peak upstream of the cyclin-dependent kinase inhibitor 3 (*CDKN3*) gene, implying that *CDKN*3 may be a direct target gene of KLF7 (Sun Y, 2016, unpublished data). Given that CDKN3 is a cell cycle regulator [23], in this study, we aimed to investigate the function of CDKN3 in preadipocyte proliferation and whether it mediates the role of KLF7 in preadipocytes. We characterized the structure of the *CDKN*3 promoter and found that KLF7 regulates *CDKN*3 expression. In addition, we explored the role and molecular mechanism of CDKN3 in preadipocyte proliferation. Our results showed that KLF7 promotes preadipocyte proliferation by activating the Akt pathway, which subsequently accelerated G1/S transition via the upregulation of CDKN3 transcription. To the best of our knowledge, this study is the first to reveal the mechanism by which KLF7 upregulates preadipocyte proliferation.

## Materials and methods

### Animals and Tissue sampling

Abdominal fat tissue samples were obtained from Arbor Acres (AA) commercial broilers (Aviagen Broiler Breeders, Beijing, China). In total, 21 birds (3 birds per time point) were slaughtered at 1–7 weeks of age. Abdominal fat tissues were collected and snap-frozen and stored in liquid nitrogen until RNA extraction.

The animal work was guided by the statute built by the Ministry of Science and Technology of the People’s Republic of China (Approval number: 2006-398) and were approved by the Laboratory Animal Management Committee of Qiqihar University.

### Cell Lines and Culture

The immortalized chicken preadipocyte ICP1 [24] and chicken fibroblast DF-1 [25] cell lines were kindly provided by the Key Laboratory of Chicken Genetics and Breeding, Ministry of Agriculture and Rural Affairs, Northeast Agricultural University (Harbin, China). ICP1 cells have been widely used to study chicken adipogenesis [26, 27], as they present the same morphological and differentiation characteristics as primary preadipocytes [24, 26, 27]. DF-1, an immortalized chicken fibroblast cell line, can be easily transiently transfected with exogenous DNA. DF-1 is often used for transcriptional regulation analysis in birds and is an ideal cell model for luciferase reporter gene assays [28-31]. ICP1 and DF-1 cells were respectively cultured in Dulbecco’s modified Eagle’s medium (DMEM)/F12 and high-glucose DMEM (Gibco, Carlsbad, CA, USA) supplemented with 10% fetal bovine serum (Biological Industries, Germany) and 1% penicillin-streptomycin solution (Beyotime, Shanghai, China), at 37 °C in the presence of 5% CO_2_.

### Plasmid Construction

The overexpression plasmid pCMV-Myc-KLF7 was constructed as described in our previous report [4]. To determine the core region of the *CDKN*3 promoter, four fragments (nucleotides [nt] −1912/–7, −758/–7, −450/–7, and −160/–7) were amplified using specific primers containing the *Kpn*I and *Hind*III restriction sites **(Fig. 2B)**. The PCR products were cloned into the pGL3-Basic (Promega, Madison, WI, USA) vector using a ClonExpress Entry One Step Cloning Kit (Vazyme, Nanjing, China). Through DNA synthesis (Genewiz, Suzhou, China), the four putative KLF7-binding sites (nt −137/–128, −87/–79, −72/–64, −56/–47) were specifically deleted in the context of pGL3-*CDKN*3 (nt −160/–7), and the resultant reporter plasmids were designated as pGL3-*CDKN*3-DM1, pGL3-*CDKN*3-DM2, pGL3-*CDKN*3-DM3, and pGL3-*CDKN*3-DM4, respectively. The full-length coding region of chicken *CDKN*3 (accession number: NM_001252162.1) was amplified by RT-PCR and cloned into the empty vector pCMV-HA (Clontech, San Francisco, CA, USA) to construct the overexpression plasmid pCMV-HA-CDKN3. All primers used are listed in **Table 1**.

Three interference fragments, si-*CDKN*3-1, si-*CDKN*3-2, and si-*CDKN*3-3, and si-NC were designed according to the *CDKN*3 coding sequence and were synthesized at Hanbio Biotechnology (Shanghai, China). The sequences are listed in **Table 2**.

### Cell Transfection

When the cell density reached 60–70%, plasmids or constructs were transfected into the cells using Invitrogen Lipofectamine^®^ 2000 reagent (Invitrogen, Carlsbad, CA, USA). Forty-eight hours after the transfection, the cells were harvested and immediately subjected to total RNA or protein isolation. Each experiment was repeated three times independently.

### Dual-Luciferase Reporter Gene Assay

Dual-luciferase reporter assays were performed using DF-1 and ICP1 cells. To characterize the *CDKN*3 promoter, cells were transfected with pGL3-*CDKN*3 or pGL3-Basic. To analyze the effect of KLF7 overexpression on *CDKN*3 promoter activity, cells were co-transfected with pGL3-*CDKN*3 and pCMV-Myc-KLF7. The cells were lysed, and promoter activities were assessed using a Dual-Luciferase Assay System (Promega, Madison, WI, USA) according to the manufacturer’s instructions. The transfection efficiency was normalized to that of the *Renilla* luciferase vector (pRL-TK). The dual-luciferase reporter assay was performed in three independent experiments with three replicates each.

### RNA Isolation and RT-qPCR

Total RNA was extracted from ICP1 cells using RNAiso Plus (Takara Bio, Dalian, China) and was reverse-transcribed to cDNA using HiScript^®^ II Q Select RT SuperMix (Vazyme, Nanjing, China). qPCRs were run using 10-μL reaction mixtures containing 1 μL of cDNA, 0.2 μL of each primer, and 5 μL of 2× ChamQ™ SYBR^®^ qPCR Master Mix (Without ROX) (Vazyme, Nanjing, China), according to the manufacturer’s instructions. The thermal cycling conditions were 95 °C for 10 min and 35 cycles of 95 °C for 15 s and 60 °C for 30 s. Relative mRNA levels were normalized to that of an endogenous reference gene (*NONO*) and were calculated using the 2^−ΔΔCt^ method. Three independent experiments with three replicates each were performed. Primer sequences are listed in **Table 1**.

### Western Blot Analysis

Forty-eight hours after transfection, cells were lysed on ice for total protein extraction. The total protein was quantified using the bicinchoninic acid assay (Beyotime, Shanghai, China), separated by 10% polyacrylamide gel electrophoresis, and transferred to a polyvinylidene fluoride membrane (Millipore, Boston, MA, USA) following the manufacturer’s instructions. After blocking with 5% skimmed milk (BD, Biosciences, Franklin Lakes, NJ, USA), the membrane was incubated with the relevant primary antibody at 4 °C overnight, then with the secondary antibody (680LT, 680 RD, dilution ratio 1:5000; LI-COR, Lincoln, NE, USA) at room temperature for 1 h, and then scanned using a two-color infrared fluorescence imaging system (Odyssey^®^, LI-COR). After washing with western primary and secondary antibody removal solution (Beyotime), the membranes were hybridized with an internal reference anti-β-actin antibody (Beyotime, AF0003, 1:1000). The primary antibodies used were anti-HA (#3724, 1:1000; CST, USA), anti-Myc (AM926, 1:1000; Beyotime), anti-KLF7 (6334-1, 1:200; Abmart, Shanghai, China), anti-Akt (AA326, 1:1000; Beyotime), anti-Akt-Ser473 (AA329, 1:1000; Beyotime), and anti-Akt-Thr308 (AA331, 1:1000; Beyotime). Three independent experiments with three replicates each were conducted.

### Cell Viability Assay

Cell viability was assessed using the Cell Counting Kit-8 (CCK-8) assay (Dojindo, Shanghai, China). ICP1 cells were cultured in 96-well plates and transfected with pCMV-HA-CDKN3 or si-*CDKN*3 to up- or downregulate *CDKN*3 expression, respectively. Six hours after transfection, which was considered time point 0, the cells were incubated with 10 μL of CCK-8 in the dark for 2 h, after which the absorbance at 450 nm was measured using a Model 680 Microplate Reader (Bio-Rad, Hercules, CA, USA). Data were recorded at 0, 12, 24, 48, and 72 h. Three independent experiments with three replicates each were conducted.

### Cell Proliferation Assay

Cell proliferation was assessed using the 5-ethynyl-2′-deoxyuridine (EdU) assay. ICP1 cells seeded in 6-well plates were cultured to 70–80% density and then transfected with the relevant siRNAs or overexpression vector. Forty-eight hours post transfection, the cells were fixed and stained using the BeyoClick™ EdU-488 Cell Proliferation Assay Kit (Beyotime). An inverted fluorescence microscope (Olympus, Beijing, China) was used to capture cells in three randomly selected fields, and the ImageJ software was used to count EdU-stained cells. The experiment was repeated three times independently.

### Cell Cycle Analysis by Flow Cytometry

Flow cytometry (Beckman Coulter, Brea, CA, USA) was used to detect the proportion of cells in each cell cycle phase using the Cell Cycle and Apoptosis Detection Kit (Beyotime) according to the manufacturer’s instructions. ICP1 cells were transfected with plasmid DNA (pCMV-HA-CDKN3, pCMV-HA) or siRNA (si-*CDKN*3 or si-NC). Forty-eight hours later, the cells were harvested and fixed in 70% ethanol at −20 °C overnight. After removing the fixation medium by centrifugation at 2,000 rpm, 500 μg of propidium iodide staining solution (Beyotime) was added to the pelleted cells. The mixture was incubated at 37 °C for 30 min. Red fluorescence was observed by flow cytometry at an excitation wavelength of 488 nm.

### Statistical and Bioinformatics Analyses

All data are shown as means ± standard deviation (SD). Means of two groups were compared using two-way Student’s t-test. *P* < 0.05 was considered statistically significant. Correlation was calculated using Pearson’s product-moment correlation. Statistical analyses were performed using the SPSS package (IBM SPSS Statistics for Windows, v.22.0, IBM, Armonk, NY, USA). Transcription factor binding sites were predicted using the JASPAR [32] online database.

## Results

### KLF7 and CDKN3 Expression are Positively Correlated in Chicken Adipose Tissue

To explore whether KLF7 is functionally associated with CDKN3 in chicken preadipocytes, we first detected *KLF*7 and *CDKN*3 expression in abdominal adipose tissues collected from AA commercial broilers of 1–7 weeks of age **(Fig. 1)** and statistically analyzed the correlation. The results showed that *CDKN*3 expression significantly declined from 1 to 6 weeks of age **(*P* < 0.05, Fig. 1)**, whereas *KLF*7 expression tended to decline from 1 to 6 weeks of age (***P* > 0.05, Fig. 1)**. Pearson’s correlation analysis showed a significant positive correlation between *KLF*7 and *CDKN*3 expression at 1–7 weeks of age **(*r* = 0.734, *P* = 0.000, *n* = 21)**.

**Figure 1.**
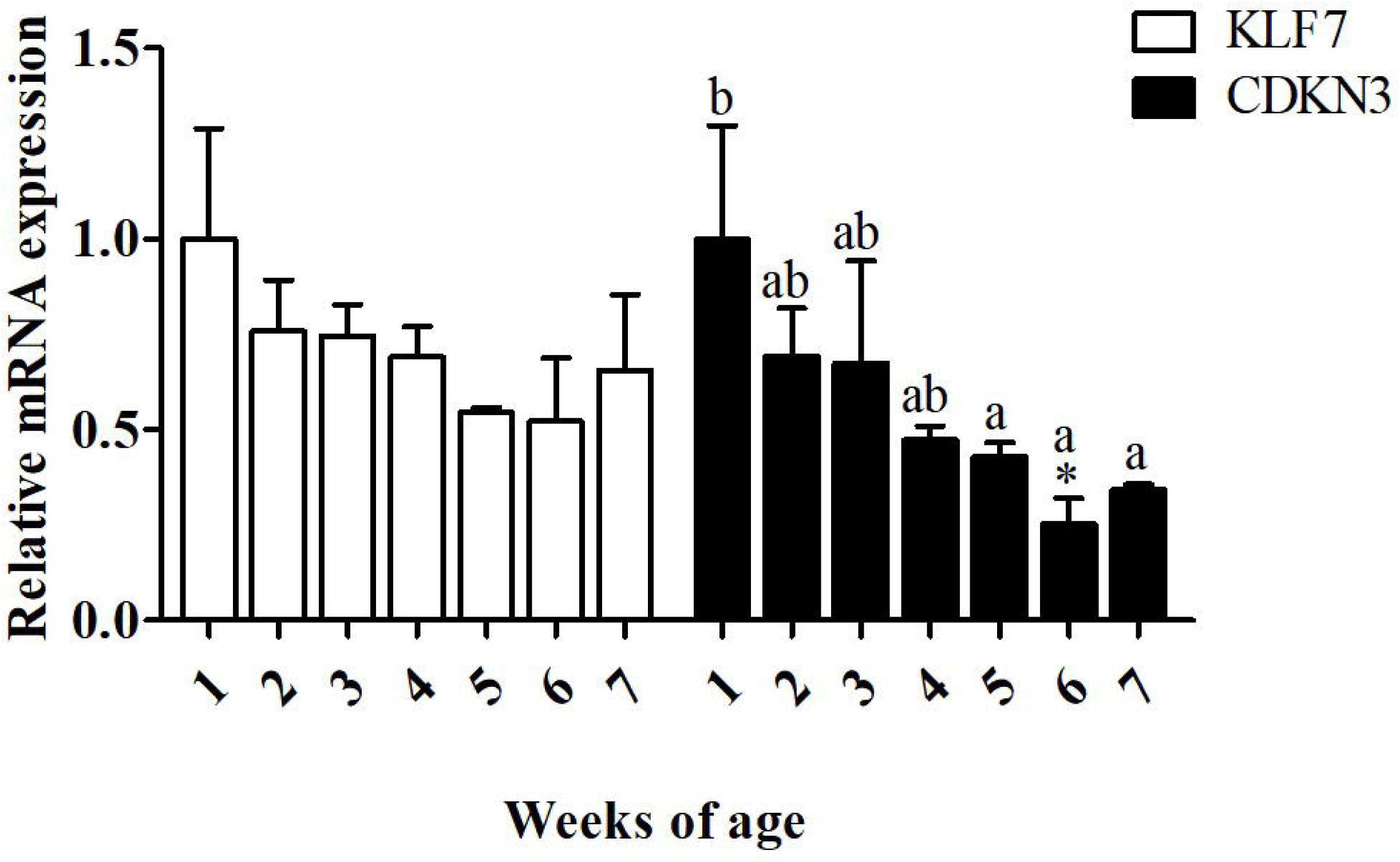
KLF7 and CDKN3 expression in abdominal adipose tissues of AA commercial broilers of 1–7 weeks of age. Gene expression was determined by RT-qPCR. Data are mean ± SD (n = 21). \**P* < 0.05, Student’s t-test.

### Characterization of the Chicken CDKN3 Promoter

To characterize the *CDKN*3 promoter, we constructed four 5′-truncated mutant luciferase reporter gene plasmids: pGL3-*CDKN*3 (nt −1912/–7), pGL3-*CDKN*3 (nt −758/–7), pGL3-*CDKN*3 (nt −450/–7), and pGL3-*CDKN*3 (nt −160/–7) **(Fig. 2A)**, and transfected them into ICP1 and DF-1 cells. Luciferase activity assay results showed that all constructs generated stronger promoter activity than empty pGL3-Basic vector **(*P* < 0.05, Fig. 2B, C)**. Notably, pGL3-*CDKN*3 (nt −160/–7) retained basal promoter activity when compared with pGL3-Basic **(*P* < 0.01, Fig. 2C, D)**. Bioinformatics analysis using the Promoter Inspector server predicted that the region nt −160/–7 contained an upstream core promoter element CGGCGCC (nt −101/–95) **(Fig. 2D)**. We also found core promoter elements (G/C)(G/C)(G/A)CGCC in this region of human, mouse, rat, pig, cow, sheep, and horse *CDKN*3 **(Fig. 2D)**. These results indicated that nt −160/–7 bp is the minimum promoter region of chicken *CDKN*3.

**Figure 2.**
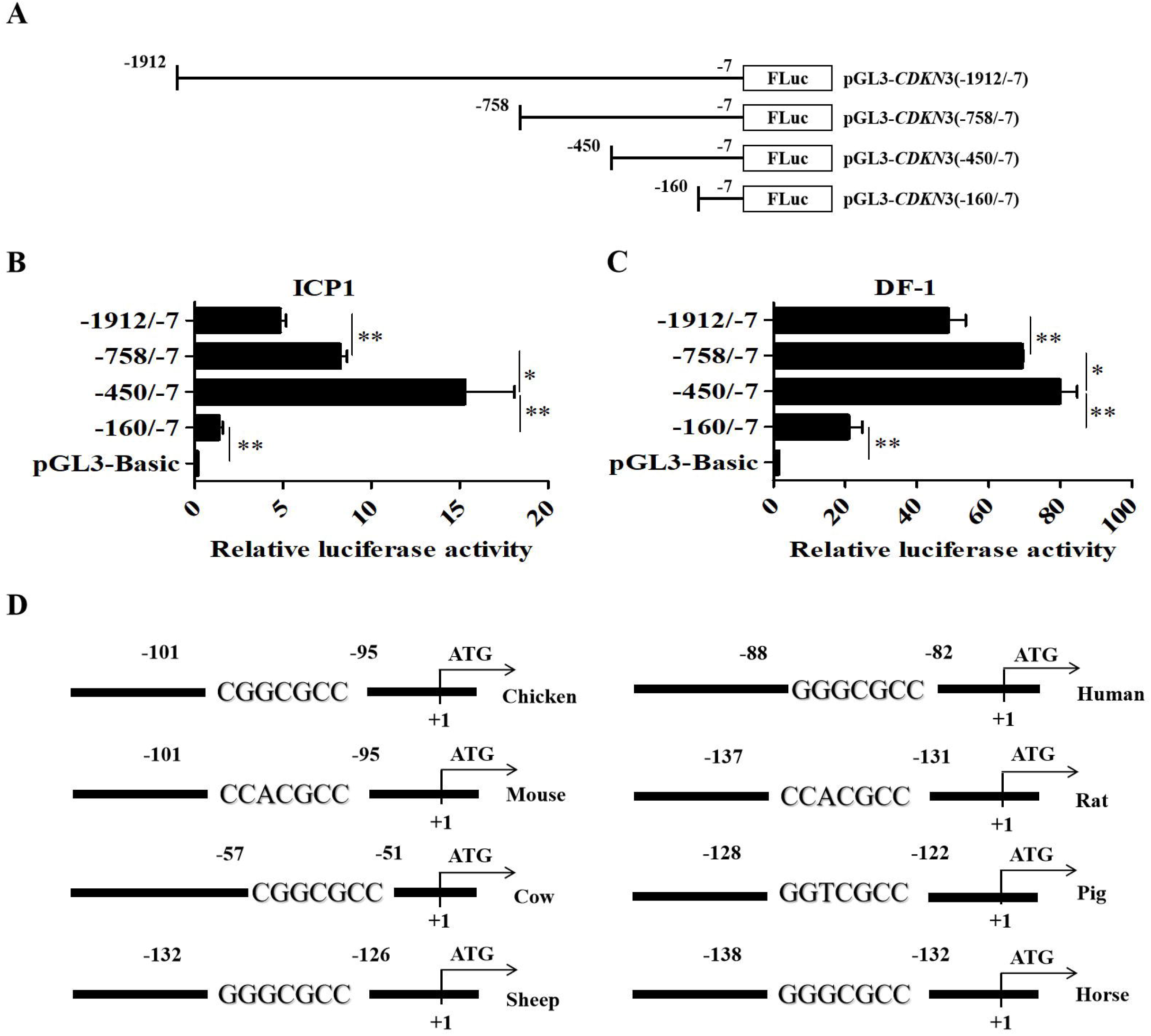
Characterization of the chicken *CDKN*3 promoter. (A) Schematic diagram of the *CDKN*3 reporter constructs used in this study. (B, C) Analysis of *CDKN*3 promoter activity. Luciferase reporter vectors were transfected into ICP1 (B) or DF-1 (C) cells with pRL-TK (transfection ratio 100:1). Relative luciferase activity was calculated as the ratio of firefly to *Renilla* luciferase activity. (D) The CGGCGCC core promoter element of chicken *CDKN*3 is located at nt −101/–95 (the ATG codon, the transcription start site of *CDKN*3, is considered position +1). Promoter core elements (G/C)(G/C)(G/A)CGCC of *CDKN*3 in various other species are shown for comparison. Data are mean ± SD. \**P* < 0.05, \*\**P* < 0.01, Student’s *t-*test.

### KLF7 Promotes CDKN3 Transcription

To determine the effect of KLF7 on promoter activity, four 5′-terminal deletion firefly luciferase reporter gene constructs were co-transfected with pCMV-Myc-KLF7 into ICP1 and DF-1 cells. Luciferase activity assay results showed that KLF7 overexpression significantly enhanced the promoter activity of all four reporter gene constructs, including the one containing the shortest *CDKN*3 promoter fragment (–160/–7) **(*P* < 0.01, Fig. 3A, B)**, suggesting that the minimal promoter, *CDKN*3 nt −160/–7, has a KLF7-binding site.

**Figure 3.**
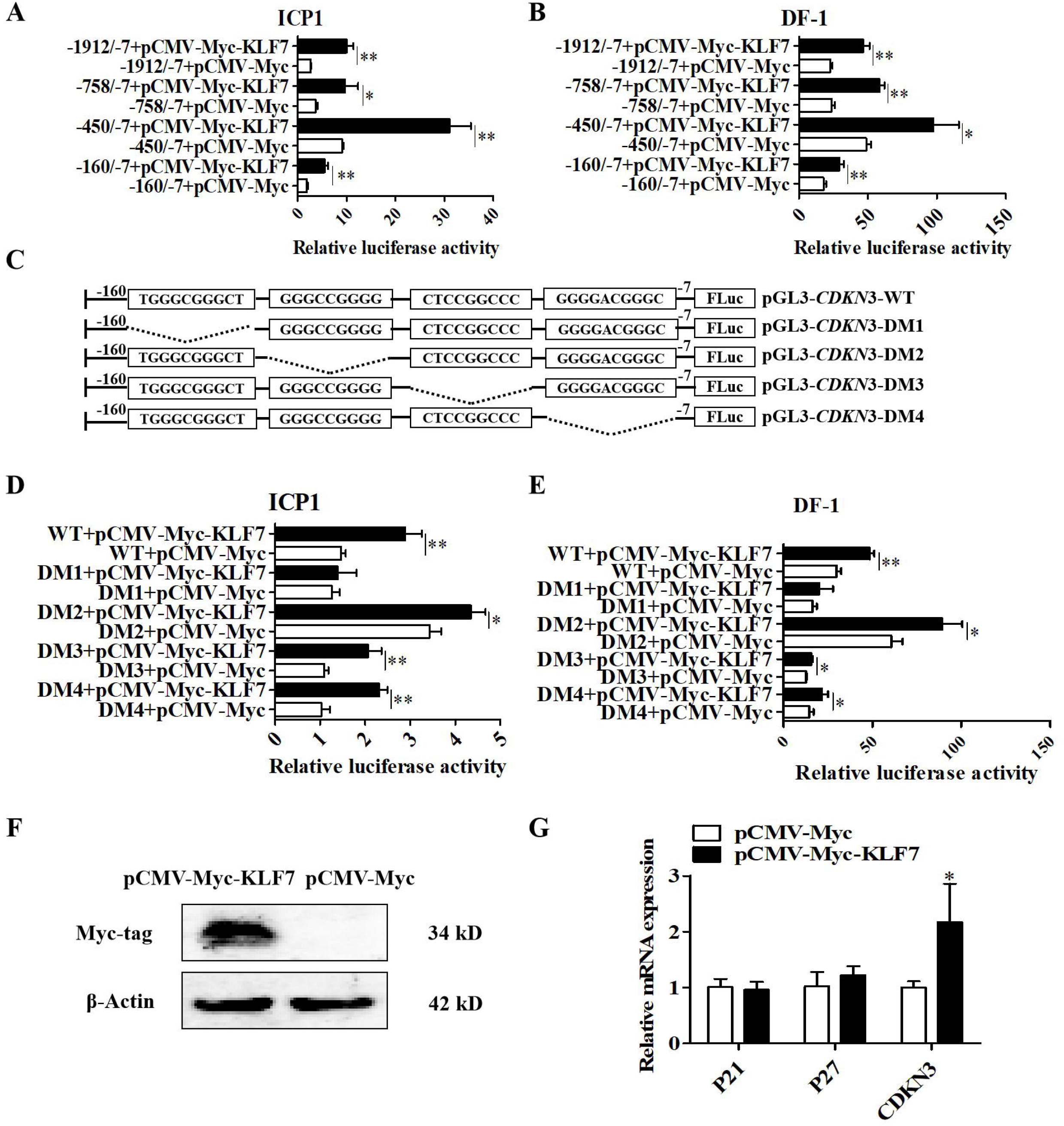
KLF7 facilitates *CDKN*3 transcription. (A, B) Reporter gene assays of the effect of KLF7 overexpression on *CDKN*3 promoter activity in ICP1 (A) and DF-1 (B) cells. Relative luciferase activity was calculated as the ratio of firefly to *Renilla* luciferase activity. (C) Schematic representation of the binding site-deletion constructs. Each of the four potential core sequences is depicted in a box. A dashed line indicates core sequence deletion. (D, E) Analysis of KLF7 binding to the *CDKN*3 promoter using a reporter gene assay. Wild-type reporter pGL3-*CDKN*3 (WT) or the DM deletion vectors and pCMV-Myc-KLF7 were co-transfected into ICP1 (D) and DF-1 (E) cells. Relative luciferase activity was calculated as the ratio of firefly to *Renilla* luciferase activity. (F) Myc protein expression levels. (G) *P*21, *P*27, and *CDKN*3 transcript levels in ICP1 cells. Data are mean ± SD. \**P* < 0.05, \*\**P* < 0.01, Student’s *t-*test.

Next, we determined the binding site of KLF7 in the *CDKN*3 promoter. We attempted to find the KLF7-binding motif revealed by our previous ChIP-seq analysis (Sun Y, 2016, unpublished data); however, unfortunately, we could not find it. Therefore, we predicted KLF7-binding sites in the *CDKN*3 promoter using the Jaspar online database [32] and found four potential KLF7-binding sites in this region **(Fig. 3C)**. We deleted the four potential binding sites based on the pGL3-*CDKN*3 (nt −160/–7) promoter construct. The deletion reporter constructs, designated pGL3-*CDKN*3-DM1, pGL3-*CDKN*3-DM2, pGL3-*CDKN*3-DM3, and pGL3-*CDKN*3-DM4, were co-transfected with pCMV-Myc-KLF7 into ICP1 and DF-1 cells. Luciferase activity assay results showed that deletion of the DM1 site TGGGCGGGCT (nt −137/–128) nearly completely abolished the effect of KLF7 on the promoter activity of pGL3-*CDKN*3 (nt −160/–7) **(*P* < 0.05)**, whereas the other three deletions had no apparent effect **(*P* > 0.05, Fig. 3D, E)**. These results indicated that the binding site TGGGCGGGCT (nt −137/–128) is required for KLF7 to exert activity on the *CDKN*3 promoter. In addition, we also statistically compared the fold-induction of the different deletion mutants to the wild-type. The results showed that the induction of DM2 and DM3 was significantly reduced in ICP1 and DF-1 cells, respectively **(*P* < 0.05)**, indicating these two sites also contribute to the response.

Studies in mammals have demonstrated that KLF7 drives neurogenesis by regulating the expression of *P*21 (also named *CDKN*1*A*) [33] and *P*27 (also named *CDKN*1*B*) [34], two other members of the CIP/KIP family. To determine whether KLF7 also acts on *P*21 and *P*27 in chicken preadipocytes, pCMV-Myc-KLF7 and pCMV-Myc were transfected into ICP1 cells, and Myc expression levels were detected using western blotting to confirm KLF7 overexpression **(Fig. 3F)**. RT-qPCR results showed that overexpression of KLF7 increased the endogenous expression of *CDKN*3 **(*P* < 0.05)**, but did not affect that of *P*21 and *P*27 **(*P* > 0.05, Fig. 3G)**.

### CDKN3 Upregulates Preadipocyte Proliferation

To detect whether CDKN3 is involved in chicken preadipocyte proliferation, *CDKN*3 expression was detected during the proliferation of ICP1 cells. A CCK-8-based cell viability assay showed that ICP1 cells gradually increased in number from 0 h to 72 h **(Fig. 4A)**, indicating that the cells were proliferating normally. RT-qPCR showed that *CDKN*3 was expressed during this process **(Fig. 4B)**, suggesting that CDKN3 plays a role in ICP1 cell proliferation.

**Figure 4.**
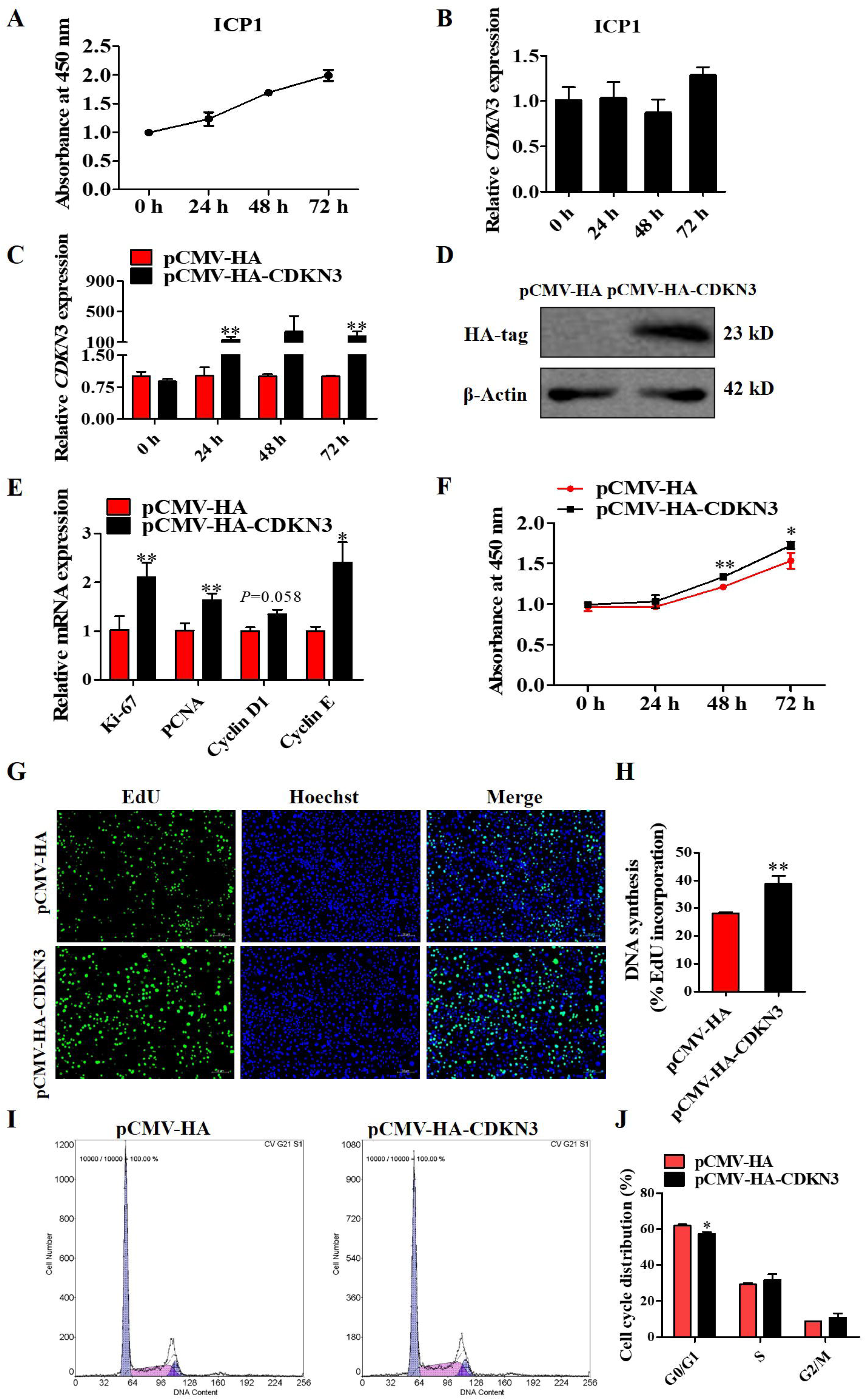
CDKN3 overexpression upregulates ICP1 cell proliferation. (A) ICP1 cell viability was determined using a CCK-8 assay. The OD_450_ value represents the proliferation capacity determined in three independent experiments. (B) RT-qPCR was used to detect *CDKN*3 expression during proliferation. (C) RT-qPCR was used to verify the overexpression of pCMV-HA-CDKN3 in ICP1 cells at 0, 24, 48, and 72 h post-transfection. (D) Western blotting was used to verify the effect of pCMV-HA-CDKN3 overexpression in ICP1 cells 48 h post transfection. Western blot analysis was repeated three times, and representative results are shown. (E) Transcript levels of *Ki-*67, *PCNA, Cyclin D*1, and *Cyclin E* were detected using RT-qPCR at 48 h post transfection. (F) The CCK-8 assay was used to detect the effect of CDKN3 overexpression on ICP1 cell proliferation. The line chart shows light absorption at 450 nm. (G, H) Proliferation of ICP1 cells transfected with pCMV-CDKN3 as assessed based on EdU incorporation (green fluorescence). (I, J) Effect of CDKN3 overexpression on the ICP1 cell cycle in ICP1 cells as measured by flow cytometry. Data are mean ± SD. \**P* < 0.05, \*\**P* < 0.01, Student’s *t-*test.

To explore the functions of CDKN3 in chicken preadipocyte proliferation, we performed overexpression and interference assays. The effects of CDKN3 overexpression **(Fig. 4C, D)** and interference **(Fig. 5A)** in ICP1 cells were analyzed using western blotting and RT-qPCR. CDKN3 overexpression promoted the expression of the proliferation markers *Ki*-67, *PCNA*, and *Cyclin E* **(*P* < 0.05, Fig. 4E)**, whereas CDKN3 interference significantly suppressed the transcript levels of these markers **(*P* < 0.05, Fig. 5B)**. Moreover, CCK-8 and EdU assay results showed that CDKN3 overexpression significantly enhanced ICP1 cell proliferation **(*P* < 0.05, Fig. 4F-H)**, whereas proliferation was significantly reduced after CDKN3 interference **(*P* < 0.05, Fig. 5C-E)**.

**Figure 5.**
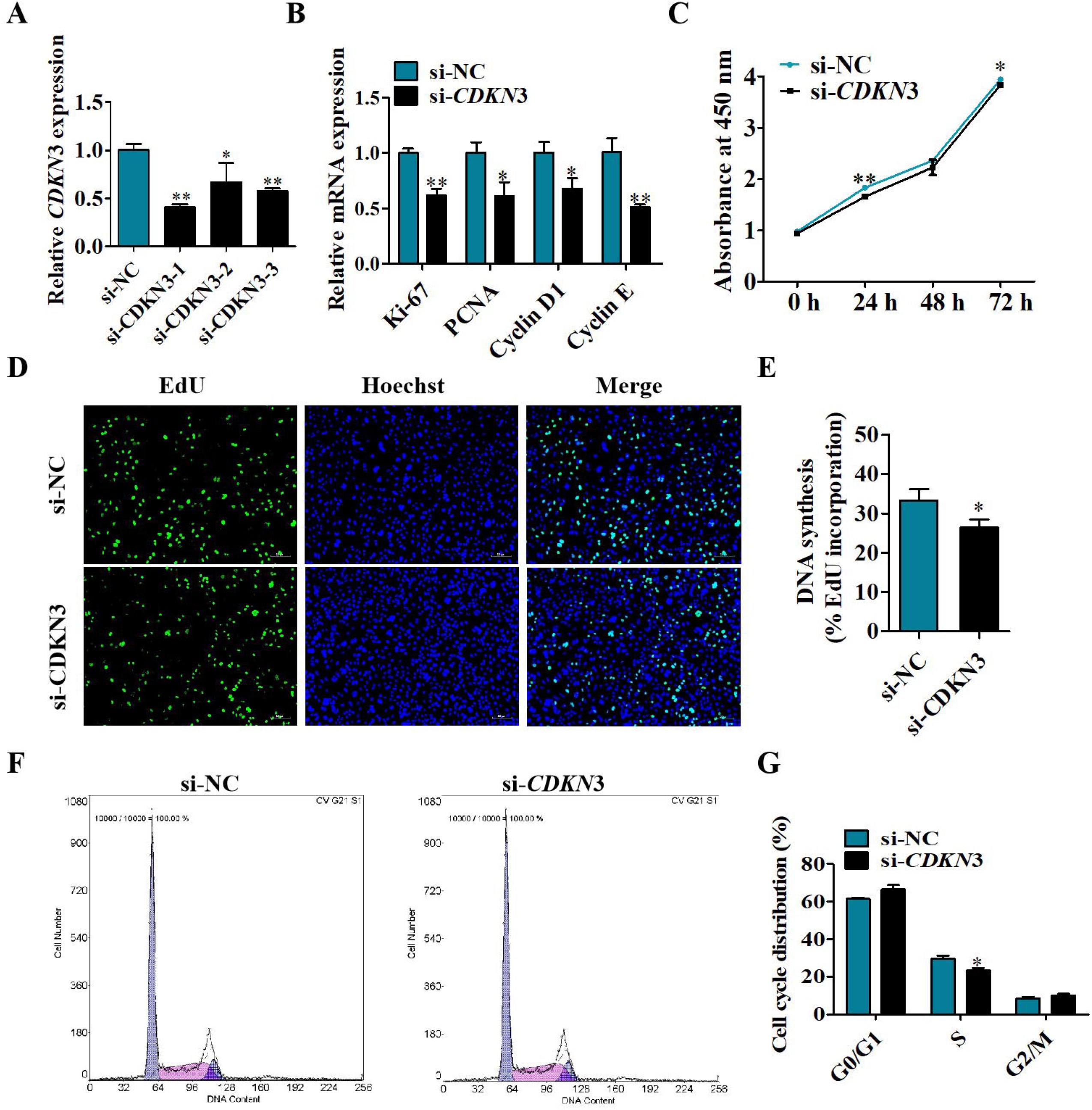
CDKN3 interference reduces ICP1 cell proliferation. (A) Effects of CDKN3 interference in ICP1 cells as detected by RT-qPCR 48 h after transfection. (B) RT-qPCR was used to detect the effect of CDKN3 interference on the expression of cell proliferation marker genes *Ki-*67, *PCNA, Cyclin D*1, and *Cyclin E*. (C) The CCK-8 assay was used to detect ICP1 cell viability upon CDKN3 interference. (D, E) Proliferation of si-*CDKN*3 transfected ICP1 cells as assessed based on EdU incorporation (green fluorescence). (F, G) Effect of CDKN3 interference on the cell cycle of ICP1 cells as assessed by flow cytometry. Data are mean ± SD. \**P* < 0.05, \*\**P* < 0.01, Student’s *t-*test.

To determine the role of CDKN3 in cell cycling, the percentage of ICP1 cells in each phase of the cell cycle was assessed using flow cytometry. Overexpression of CDKN3 significantly reduced the number of cells in the G0/G1 phase **(*P* < 0.05 Fig. 4I, J)**, whereas CDKN3 interference remarkably decreased the number of cells remaining in the S phase **(*P* < 0.05, Fig. 5F, G)**. These results implied that CDKN3 promotes the preadipocyte proliferation by driving the G1/S transition.

### KLF7 Activates the Akt Signaling Pathway by Upregulating CDKN3 Expression

KLF7 is a positive regulator of chicken preadipocyte proliferation [4]. We showed that CDKN3 promoted preadipocyte proliferation **(Fig. 4 and Fig. 5)** and *CDKN3* transcription was enhanced by KLF7 **(Fig. 3)**. Therefore, we investigated whether KLF7 promotes preadipocyte proliferation by upregulating *CDKN*3 expression using the EdU assay. The proliferation of cells transfected with pCMV-Myc-KLF7 was significantly higher than that of cells transfected with pCMV-Myc **(*P* < 0.01, Fig. 6A, B)**, indicating that KLF7 overexpression promotes preadipocyte proliferation. The proliferation of cells transfected with si-*CDKN*3 was lower than that of cells in the NC group **(*P* < 0.05, Fig. 6A, B)**, indicating that CDKN3 inference reduces chicken preadipocyte proliferation. There were no remarkable differences in proliferation between the si-NC group and KLF7 overexpression + si-CDKN3 group **(*P* > 0.05, Fig. 6A, B)**. These results suggested that KLF7 promotes proliferation by upregulating *CDKN*3 expression.

**Figure 6.**
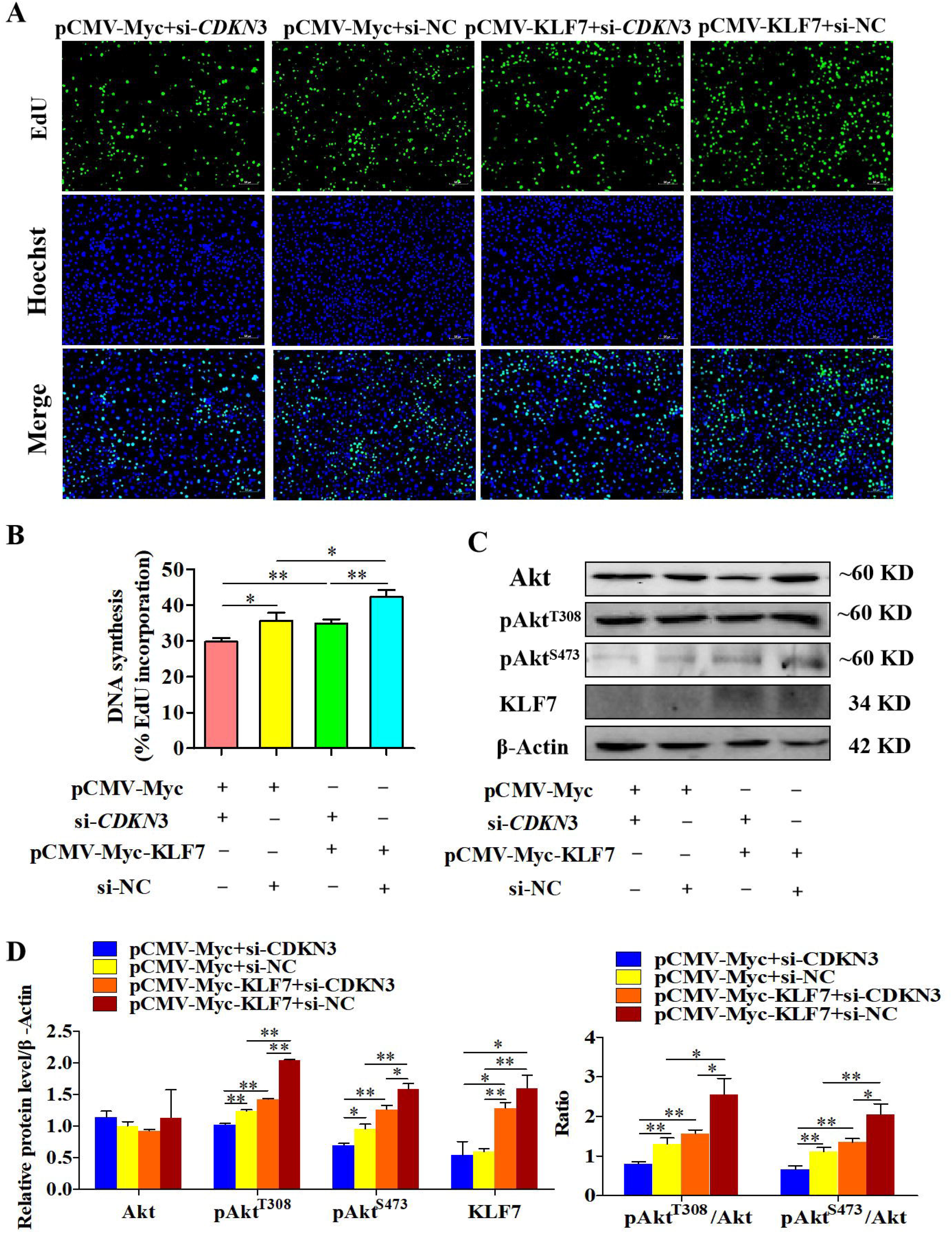
KLF7 activates the Akt signaling pathway via CDKN3. (A, B) Forty-eight hours post transfection, the proliferation of ICP1 cells transfected with pCMV-Myc-KLF7 and si-*CDKN*3 was assessed based on EdU incorporation (green fluorescence). (C, D) Western blotting was used to detect the effects of KLF7 overexpression and *CDKN*3 interference on Akt, pAkt^T308^, and pAkt^S473^ levels. The analysis was repeated three times independently, and representative results are shown. Data aremean ± SD. \**P* < 0.05, \*\**P* < 0.01, Student’s *t-*test.

As the Akt signaling pathway plays a key role in cell cycle regulation [35] and CDKN3 reportedly activates Akt phosphorylation in cancer cells [36-38], we investigated whether KLF7 activates the Akt pathway by upregulating CDKN3 in ICP1 cells. Western blot results showed that, compared with the Myc+si-CDKN3 and Myc+si-NC groups, the KLF7+si-CDKN3 and KLF7+si-NC groups showed significantly higher KLF7 expression **(*P* < 0.05)**, which demonstrated that KLF7 was successfully overexpressed. The western blot results also showed that KLF7 overexpression significantly enhanced the levels of P-Akt^Thr308^ **(*P* < 0.05)** and P-Akt^ser473^ **(*P* < 0.05)** compared to those in the control group, suggesting that KLF7 promotes proliferation by enhancing Akt signaling activation in chicken preadipocytes. The levels of P-Akt^Thr308^ **(*P* < 0.01)** and P-Akt^ser473^ **(*P* < 0.01)** were lower in the si-*CDKN*3 group than in the si-NC group, implying that CDKN3 knockdown repressed Akt signaling activation. There were no significant differences in P-Akt^Thr308^ and P-Akt^ser473^ levels between the si-NC and KLF7+si-CDKN3 groups **(*P* > 0.05, Fig. 6C, D)**. Together, the present findings revealed that KLF7 overexpression facilitates Akt pathway activation by upregulating CDKN3.

## Discussion

KLF7 is a positive regulator of chicken preadipocyte proliferation [4]; however, the underlying mechanism remained poorly understood. The current study confirmed that KLF7 overexpression promoted preadipocyte proliferation (**Fig. 6A, B**) and showed that this promotion is achieved via the Akt signaling pathway **(Fig. 6C, D)**. Further, we found that KLF7 upregulated *CDKN*3 promoter activity, and the binding site TGGGCGGGCT (–137/–128 nt; DM1) is required for KLF7 to exert activity on the *CDKN*3 promoter **(Fig. 3D, E)**. In addition, KLF7 overexpression promoted the endogenous expression of *CDKN*3 **(Fig. 3G)**. Building upon our previous ChIP-qPCR and ChIP-seq-based finding that KLF7 can bind to the promoter region of *CDKN*3, we here demonstrated that DM1 likely mediates KLF7 binding to the *CDKN*3 promoter to drive *CDKN*3 transcription. However, as we did not assess combined binding site mutations, and the fold-induction of DM2 and DM3 was significantly reduced compared with the wild-type, we cannot assert that DM1 is the only binding site. Moreover, deletion of the binding site may have led to the formation of new binding sites that affected *CDKN*3 promoter activity. Despite these limitations, our results still revealed the mechanism by which KLF7 facilitates preadipocyte proliferation.

P21 (also named CDKN1A) [33] and P27 (also named CDKN1B) [34] are CIP/KIP family members [39, 40]. P21 can bind to CDK to inhibit its interaction with substrates such as Rb family members, ultimately repressing G1/S progression [41]. P27 also inhibits CDK activity, thus blocking G1/S transition and cell cycle progression [42]. KLF7 reportedly targets *P*21 and *P*27 in various mammalian cells. In nerve cells, KLF7 promotes *P*21 and *P*27 expression to drive axon outgrowth [33, 34]. KLF7 also maintains muscle stem satellite cells in quiescence and drives cell cycle exit by acting on *P*21 [43]. The role of KLF7 in preadipocyte proliferation in mice remains unclear. Interestingly, unlike murine *KLF*7, chicken *KLF*7 upregulated CDKN3 expression, but had no effect on *P*21 and *P*27 in preadipocytes **(Fig. 3G)**, implying that the molecular mechanisms of KLF7 in preadipocytes differ between birds and mammals. In the present study, we performed gene expression analysis of chicken *CDKN*3, but due to lack of available chicken antibodies, we did not perform the protein expression analysis, it is worth performing the protein expression analysis. In addition, we did not test the effects of KLF7 knockdown. For some reason, the chemically synthesized KLF7 siRNAs did not work. Thus, it would be worth further exploring the functions of chicken KLF7 and CDKN3 *in vivo* or *in vitro* using CRISPR/Cas9 assays.

Preadipocyte proliferation is controlled by a complex network of transcription factors [4-6]. For instance, BMP4 upregulates the proliferation of ICP1 cells by promoting the G1/S transition [44], whereas ALDH1A1 is a negative regulator of preadipocyte proliferation and inhibits the G1/S transition [45]. Many other factors, such as PPARγ [46], RB1 [47], HOPX [48], and TCF21 [49], also play negative regulatory roles in preadipocyte proliferation. In the current study, we showed for the first time that CDKN3 promotes preadipocyte proliferation, as evidenced and validated by three independent methods (CCK-8 and EdU assays, and flow cytometry, **Figs. 4 and 5**), that were repeated at least three times independently. Our results extend our knowledge of the molecular mechanism underlying chicken preadipocyte proliferation. Because the siRNA (si-*CDKN*3) we used acted for a short time, or may have been at the limit of transfection efficiency, the CCK-8 results showed a small difference. It may be worth further exploring the functions of CDKN3 using the CRISPR/Cas9 assay or by constructing lentiviral vectors.

CDKN3, also known as CDI1, CIP2, KAP, or KAP1, was first discovered as a binding protein of cyclin-dependent kinase 2 (CDK2). Studies have shown that CDKN3 promotes the proliferation of breast [50] and prostate [51] cancer cells. Consistent here with, we found that CDKN3 induced preadipocyte proliferation by driving the G1/S transition and inducing the expression of cyclin E, a G1/S progression promoter. Interestingly, our data contradicted a previous finding that CDKN3 inhibits the G1/S transition by dephosphorylating CDC2^Thr161^ in brain glioma cells [52]. This discrepancy suggests that CDKN3 drives or represses cell proliferation in a cell- or context-specific manner.

The Akt/mTOR pathway plays critical roles in regulating the cell cycle and apoptosis [53]. Several studies have reported that Akt pathway activation promotes preadipocyte proliferation [54, 55]. We found that CDKN3 increased Akt pathway activation in preadipocytes **(Fig. 6C, D)** and promoted preadipocyte proliferation (**Figs. 4, 5, and 6C, D**). Moreover, KLF7 induced Akt signaling activation by targeting CDKN3 **(Fig. 7)**. CDKN3 promotes the ubiquitination and subsequent phosphorylation of Akt by preventing the interaction between SKP2 and CDK2 via the dephosphorylation of CDK2 [37]. The Akt pathway also has important functions in preadipocyte differentiation. Studies have shown that Akt pathway activation promotes the expression of PPARγ and C/EBPα, two key regulators of differentiation, thus accelerating preadipocyte differentiation [56-58]. Of note, KLF7 downregulates preadipocyte differentiation [4]. However, our results indicated that both KLF7 and CDKN3 accelerated Akt pathway activation in preadipocytes. Altogether, these data suggest that KLF7 boosts preadipocyte proliferation by promoting Akt signaling via targeting CDKN3, whereas in preadipocyte differentiation, KLF7 may not play an inhibitory role via CDKN3 and Akt pathway activation.

**Figure 7.**
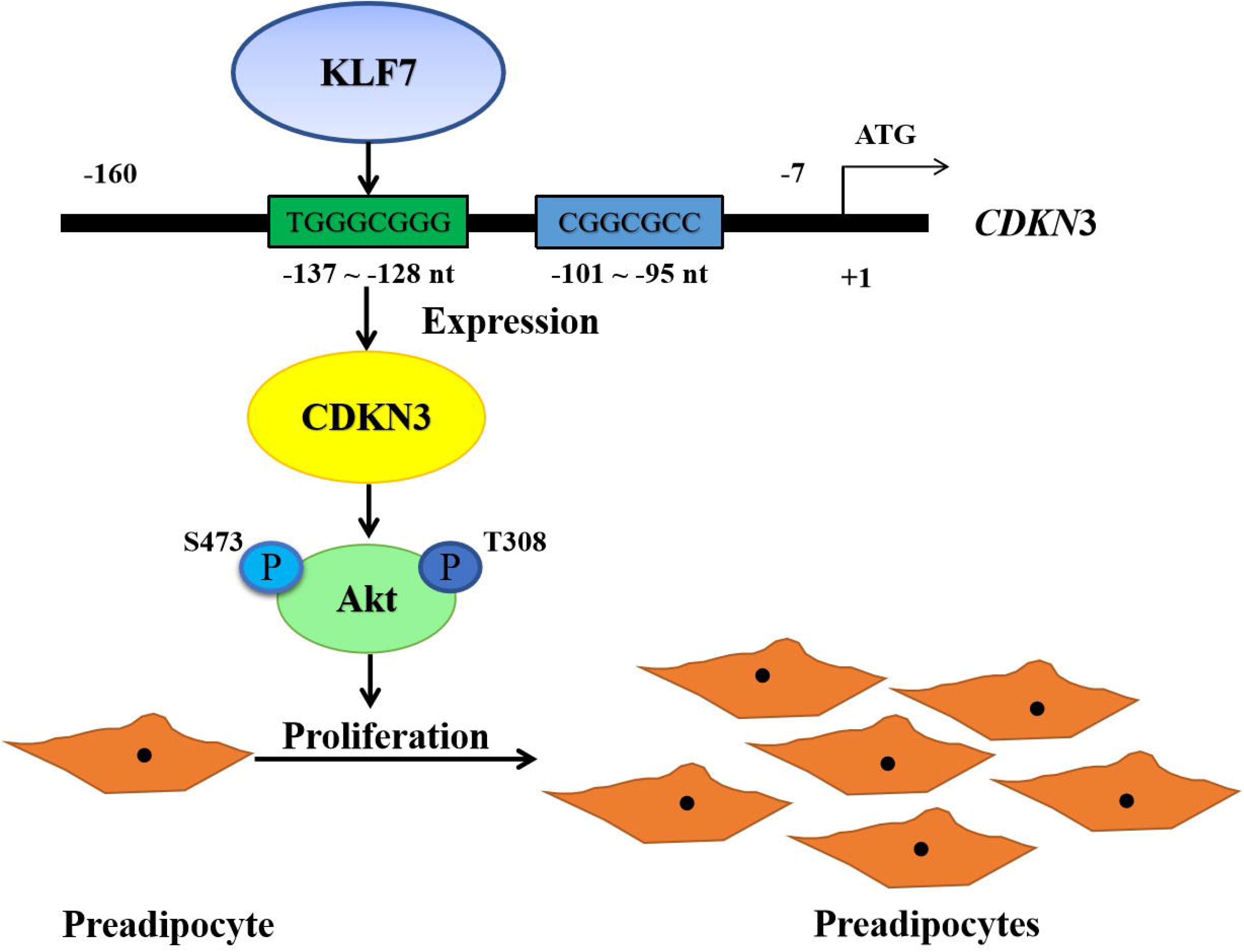
KLF7 promotes the proliferation of preadipocytes via CDKN3-mediated regulation of Akt phosphorylation. The transcription factor KLF7 can directly bind to the promoter region of *CDKN*3 to promote *CDKN*3 expression. CDKN3 drives the G1/S transition, thus promoting ICP1 cell proliferation by activating the Akt pathway via Ser and Thr phosphorylation.

Chicken is an ideal model species for studying adipogenesis and adipose biology in view of their natural hyperglycemia, insulin resistance, hepatic fatty acid synthesis, and reproductive system [22]. Especially, their high glycemia and low sensitivity to exogenous insulin (particularly in adipose tissues) make them a relevant model for studies on human obesity, insulin resistance, and type 2 diabetes [20, 21, 59]. In this study, we showed that KLF7 increased the number of ICP1 cells by promoting CDKN3 transcription. These results provide insights into the molecular regulatory network of preadipocyte proliferation and contribute to understanding human adipogenesis and obesity, which is of great significance given the high prevalence of obesity nowadays. In addition, our findings suggest that CDKN3 may be a potential therapeutic target for human obesity and obesity-related diseases.

In conclusion, we showed for the first time that *CDKN*3 is a target gene of the transcription factor KLF7 in preadipocytes. CDKN3 promotes preadipocyte proliferation by driving the G1/S transition. By upregulating CDKN3, KLF7 accelerates Akt signaling pathway activation, thereby facilitating preadipocyte proliferation in chickens.

## Acknowledgements

We would like to thank the Key Laboratory of Chicken Genetics and Breeding, Ministry of Agriculture and Rural Affairs, Northeast Agricultural University, for kindly providing chicken immortalized chicken preadipocyte cell line **(ICP1)**. We would like to thank Editage (www.editage.cn) for English language editing.

