## Supplementary material for "KLF7 Promotes Preadipocyte Proliferation via Activation of the Akt Signaling Pathway by Cis-regulating CDKN3": Table 1

### Table 1. Primer sequences

| Gene name | Primer sequence (5´-3´) | GenBank accession |
|---|---|---|
| *CDKN*3 −1912/−7 | F: atttctctatcgata ggtacc TAAGGCAATCACAGCACAA<br>R: cagtaccggaatgcc aagctt GGCTGCGGATGGAGAAGA | NC_052536.1 |
| *CDKN*3 −758/−7 | F: atttctctatcgata ggtacc CCGGAACGAAAGGCTGAG<br>R: cagtaccggaatgcc aagctt GGCTGCGGATGGAGAAGA | NC_052536.1 |
| *CDKN*3 −450/−7 | F: atttctctatcgata ggtacc CAGTGCTCCTCATCCCA<br>R: cagtaccggaatgcc aagctt GGCTGCGGATGGAGAAGA | NC_052536.1 |
| *CDKN*3 −160/−7 | F: atttctctatcgata ggtacc CTCGACCAATGAGATGCG<br>R: cagtaccggaatgcc aagctt GGCTGCGGATGGAGAAGA | NC_052536.1 |
| *CDKN*3 | F: tggccatggaggccc gaattc TGCACGTAGCCGGCTTTGACTC<br>R: gcggccgcggtac ctcgag TGCAGCACGAGTTCACCGTGA | NM_001252162.1 |
| RT-qPCR *CDKN*3 | F: TGTCCTGGCTCCCTCTGTC<br>R: CATCCTTAAACCTGCAACCC | NM_001252162.1 |
| RT-qPCR *CDKN*1*A* | F: CCCAGGGATGGTGTCG<br>R: GGGCTTATCGTGGACAACT | NM_204396.1 |
| RT-qPCR *CDKN*1*B* | F: CGCTTCTCAGGGAATCTCA<br>R: GAAACCTCCTCTTCTGTTGTG | NM_204256.2 |
| RT-qPCR *Ki*-67 | F: AGGTCCGTTCCCTCGTT<br>R: CATTGTCGTCTGGGTCATC | XM_422088.5 |
| RT-qPCR *PCNA* | F: GTGCTGGGACCTGGGTT<br>R: CGTATCCGCATTGTCTTCT | NM_204170.2 |
| RT-qPCR *Cyclin D*1 | F: AGAAGTGCGAAGAGGAAGT<br>R: TGATGGAGTTGTCGGTGTA | NM_205381.1 |
| RT-qPCR *Cyclin E* | F: TTTGCTATGGCTATAAGGG<br>R: TGTGGTGGCGTAAGGA | NM_001031358.1 |
| RT-PqCR *NONO* | F: AGAAGCAGCAGCAAGAAC<br>R: TCCTCCATCCTCCTCAGT | NM_001031532.1 |
