## Supplementary material for "KLF7 Promotes Preadipocyte Proliferation via Activation of the Akt Signaling Pathway by Cis-regulating CDKN3": Table 2

### Table 2. siRNAs used in this study

| Name | Sequence (5´-3´) |
|---|---|
| si-*CDKN*3-1 | F: AGACAAUUAUCCACUGUUAdTdT |
| | R: UAACAGUGGAUAAUUGUCUdTdT |
| si-*CDKN*3-2 | F: GUUUGUGCUUUGUACCAAAdTdT |
| | R: UUUGGUACAAAGCACAAACdTdT |
| si-*CDKN*3-3 | F: CGGAUGUAGUGACACCUCAdTdT |
| | R: UGAGGUGUCACUACAUCCGdTdT |
| si-NC | F: UUCUCCGAACGUGUCACGUdTdT |
| | R: ACGUGACACGUUCGGAGAAdTdT |
